# Determining the Depth Limit of Bioluminescent Sources in Scattering Media

**DOI:** 10.1101/2020.04.21.044982

**Authors:** Ankit Raghuram, Fan Ye, Jesse K. Adams, Nathan Shaner, Jacob T. Robinson, Ashok Veeraraghavan

## Abstract

Bioluminescence has several potential advantages compared to fluorescence microscopy for *in vivo* biological imaging. Because bioluminescence does not require excitation light, imaging can be performed for extended periods of time without phototoxicity or photobleaching, and optical systems can be smaller, simpler, and lighter. Eliminating the need for excitation light may also affect how deeply one can image in scattering biological tissue, but the imaging depth limits for bioluminescence have yet to be reported. Here, we perform a theoretical study of the depth limits of bioluminescence microscopy and find that cellular resolution imaging should be possible at a depth of 5-10 mean free paths (MFPs). This limit is deeper than the depth limit for confocal microscopy and slightly lower than the imaging limit expected for two-photon microscopy under similar conditions. We also validate our predictions experimentally using tissue phantoms. Overall we show that with advancements in the brightness of bioluminescent indicators, it should be possible to achieve deep, long-term imaging in biological tissue with cellular resolution.

## Introduction

The imaging depth in biological tissue is typically limited by scattering; on average, a photon scatters more than ten times within the first millimeter of most biological tissues [19,20]. This scattering results in blurring of the image and reduced image contrast. Together these effects limit the maximum depth at which cellular-resolution imaging is possible [19].

To improve image quality within scattering tissue, researchers often perform in *vivo* fluorescence imaging using confocal, two-photon (2P), or three-photon (3P) microscopy [8,23]. These techniques help to reduce out-of-plane fluorescence, which improves image contrast, but they rely on scanning laser systems. These scanning laser systems are not only complex but subject the tissue to high intensities that lead to photobleaching and phototoxicity, limiting the duration of an imaging experiment [31,32].

Bioluminescence provides a promising alternative technique that eliminates the need for excitation light, making it possible to perform imaging experiments for extended periods of time with simplified, light-weight imaging systems [16,30]. Bioluminescence is a form of chemiluminescence that generates light through a chemical reaction without the need for excitation light. In this way, bioluminescent proteins act as sources of light within the tissue, avoiding harmful side effects of excitation light like photobleaching and phototoxicity [16,30]. Additionally, light collection for bioluminescent imaging does not require a light source, color filters, or dichroic mirrors allowing wide-field microscopes [30,32] or their lensless counterparts [24] to be simpler and smaller than fluorescent systems.

A less-discussed advantage of bioluminescence is the fact it may be possible to image more deeply into a scattering biological sample than is possible with epifluorescence [30,32]. Because bioluminescence does not require excitation light, imaging contrast does not suffer from autofluorescence and excitation scattering, which are the primary factors that limit deep fluorescent imaging in scattering media [20]. While bioluminescence imaging still suffers from attenuation and blurring due to emission scattering, with a bright enough source, one would expect to achieve a greater imaging depth than epifluorescence [7,26,34].

In this paper, we quantitatively address the relationship between the bioluminescence brightness and imaging depth in scattering media. Using a combination of Monte Carlo simulation and experiments in tissue phantoms, we find the following: (1) The depth limit for bioluminescent imaging ranges from 5-10 MFPs. (2) As the brightness of bioluminescent reporters increases, imaging depth and spatial resolution also increase. (3) With the available bright bioluminescent indicators like eNanoLantern [26,34], we expect that a 2 *nM* concentration of reporters is needed to achieve the necessary signal to image through > 5 MFPs of scattering media.

### Fluorescence Imaging

Fluorophores emit light that is red-shifted compared to the light that they absorb, allowing one to use color filters to isolate light that is emitted by the scene [5, 23]. Fluorescence can be used to image neural activity by monitoring membrane voltage or cellular Ca^2^ + concentration [13,25]. In recent years, genetically encoded Ca^2+^ indicators (GECIs) have shown promise as *in vivo* indicators of neuronal activity because they can respond to changes in cytosolic Ca^2+^ brought about by opening and closing of Ca^2+^ channels during an action potential and can be expressed in specific neuronal populations [14]. To record neural activity from cells below the cortical surface, one must image through several hundred microns into scattering brain tissue. Due to excitation and emission scattering, signal originating from these deeper regions profoundly diminishes. To sufficiently illuminate deeper portions of tissue in standard epifluorescence microscopy, increased excitation energy is used to obtain a larger emitted signal [5]. However, out-of-plane autofluorescence and excitation scattering also increase and reduce contrast [30], limiting the depth limit to 3 MFPs for epifluorescence imaging [20].

Confocal and 2P microscopes were developed to combat these problems by reducing the amount of detected background light [2,28]. Confocal microscopy introduces a pinhole at the illumination and imaging plane in order to image a single scene point at a time, blocking out out-of-plane fluorescence (a combination of autofluorescence and out-of-plane probe fluorescence). The scene point is scanned across the sample to create the full image [28]. 2P microscopy relies on a nonlinear excitation process where fluorescence intensity scales as the square of intensity, making it possible to better restrict fluorescence to the focus of the laser spot. Due to the extremely low probability of excitation outside of the focal spot, 2P microscopy does not require a pinhole to reject out-of-plane fluorescence [2]. However, these methods, along with conventional fluorescence imaging, suffer from probe photobleaching and excitation light-induced phototoxicity, making long term studies challenging [30, 32]. Also, in the cases of confocal and 2-photon microscopy, a high powered laser and complex optical systems are needed to perform real-time imaging. These systems can be relatively large and are particularly challenging to implement in a small form-factor design for in vivo imaging.

### Bioluminescence: Potential and Pitfalls

Bioluminescence is a form of chemiluminescence that emits light from chemical reactions. Bioluminescent proteins produce light via oxidation of a small-molecule luciferin by a luciferase or photoprotein. Luciferases display constitutive activity, while photoproteins, like aequorin, produce light in a Ca^2+^-dependent manner [18]. Aequorin was first isolated from jellyfish in 1962 [27], and since then, dozens of other bioluminescent proteins have been cloned and engineered into more catalytically active and more spectrally diverse bioluminescent sources [31]. Beyond improvements in the enzymes themselves, the bioluminescence quantum yield of luciferases can be amplified through Forster resonance energy transfer (FRET) to a fluorescent protein. Among the first demonstrations of this principle was the creation of “NanoLantern,” in which *Renilla* luciferase acts as a FRET donor to the yellow fluorescent protein Venus, resulting in a large increase in emitted photons. Scientists are already using proteins like NanoLantern (and variants) for tissue imaging. For example, the NanoLantern-derived calcium sensor CalfluxVTN can be expressed via viral transduction, along with optogenetic actuators, to manipulate and monitor neural activity in rat hippocampus. [26, 34].

Bioluminescence holds a number of benefits over fluorescence imaging techniques such as increased signal-to-noise ratio, easy implementation in a small form-factor microscope since it does not need any expensive equipment, and no harmful biological effects due to excitation light. All three of these benefits point to long-term, deep-tissue, in-vivo experiments; a problem that is hard to address with current fluorescence imaging techniques. The main drawback for bioluminescence microscopy is low signal amplitude, however ongoing research is aimed at increasing the brightness of these indicators.

## 1 Computational Modeling for Bioluminescence Microscopy

Our goal in this paper is to characterize and understand the imaging performance that can be achieved using bioluminescent microscopy. To this end, we develop a computational model that allows us to simulate how the properties of the scattering tissue and the bioluminescent probes affect image quality.

This computational model has three major advantages. Firstly, it allows us to estimate how improvements in bioluminescence technology, such as brighter bioluminescent probes, will affect image quality. Secondly, this computational model allows us to systematically vary the relevant system parameters such as probe brightness, tissue scattering coefficient, depth of imaging, sparsity (or density) of sources in the scene, etc., to explore imaging performance over a large potential design space. Finally, we can use the computational model to predict the performance of the current state-of-art bioluminescent microscopy.

### 1.1 Computational Model for Bioluminescence

Light propagation through scattering media can be accurately modeled using the radiative transfer equation (RTE). The standard integro-differential equation for a plane-parallel medium is defined as [20]:

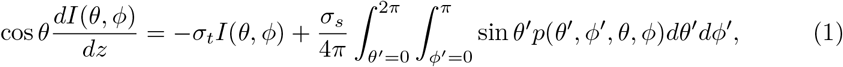

where *θ* and *φ* are the azimuthal and elevation angles, respectively. *I* is the intensity at a given depth *z*. p(.) is the probability that a photon traveling in the (*θ*’, φ’) direction scatters into the (*θ*, φ) direction. *σ_t_* corresponds to the extinction cross section, made up of *σ_s_*, scattering cross section, and *σ_a_*, absorption cross section (not shown). For situations in which there is little to no absorption, the extinction cross-section can be represented with just the scattering cross-section (*σ_t_* = σ_s_), which is the case for brain imaging [20], as most brain tissue is predominantly scattering. The change in the intensity of light at a certain (*θ*, φ) with respect to depth in the medium (left-hand side) depends on two processes (if there is no intensity generation at this *z*): loss of intensity that is either absorbed or scattered (right hand side, term 1) or light from another direction (*θ*’,φ’) scattering into the (*θ*, φ) direction (right hand side, term 2).

Currently, there are no analytical solutions to the RTE. As a consequence, Monte Carlo simulations have been used to estimate the solution to the RTE by converting the deterministic radiative-transfer process into a probabilistic process [17,33]. By running many samples, an accurate estimate is achieved. We follow this line of work and employ Monte Carlo modeling to study and characterize light propagation.

Figure 1a shows a diagram of the model to render different instantiations of bioluminescent sources. Virtual photons originated at each of the 10*μm* × 10*μm* × 10*μm* sources. A feature size of 10*μm* was chosen because it closely resembles the size of small neurons [4]. Sources emitted photons isotropically, which interacted with the scattering medium. The arrival of scattering molecules to the photon was modeled as a Poisson process. The inter-arrival time of scattering molecules (the distance the photon traveled before scattering) is given by *l* = *l_s_* log ξ where *l_s_* is the mean free path (MFP) length defined as the average distance a photon travels before scattering within a medium and ξ ~ *U*[0,1]. Once a photon reached a scattering event, a new scattering direction (given by the Henyey-Greenstein phase function) is computed [20]:

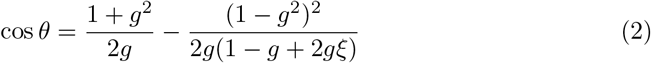

**Figure 1.**
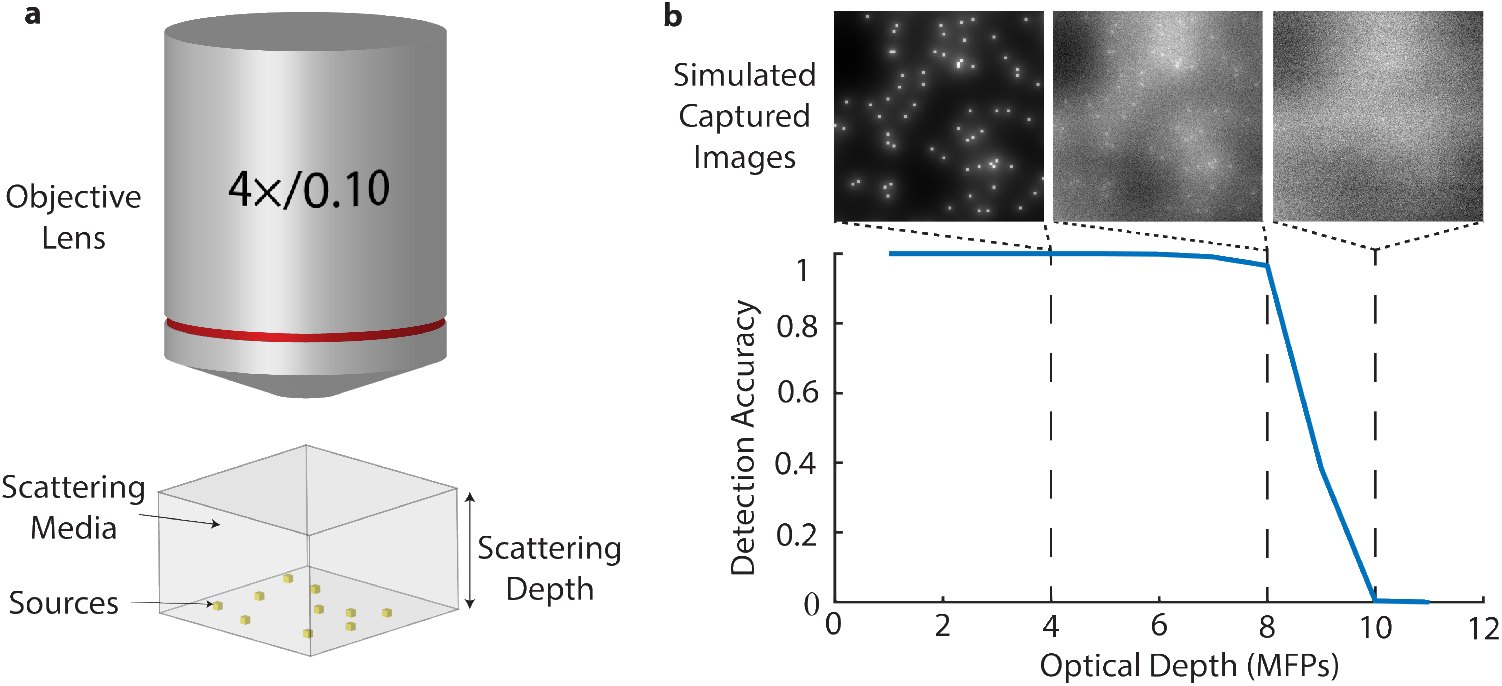
The depth limit of bioluminescent sources in scattering media. (**a**) shows the diagram of the Monte Carlo simulation. Photons originate from sources on one side of the scattering medium and travel to the imaging system represented by the 4× objective to form an image. (**b**) show the resultant images formed by running the Monte Carlo simulation with sources starting at different scattering depths and the corresponding detection accuracy at those depths. Detection accuracy is the percent of sources correctly detected, given a fixed false discovery rate (0.05). The source density and emission rate per source was 131 sources/mm^2^ with 10^10^ photons/source, respectively.

Here, *g* is the anisotropy parameter which is about 0.9 for brain tissue [12]. After scattering, the azimuth angle was uniformly sampled to determine the new scattering direction of the photon. This Monte Carlo model was adapted from Pediredla *et al.*’s simulation for single plane illumination microscopy [20]. Photons that reached the end of the scattering tissue were traced onto the objective lens of a 4*f* system. To model the lens, we used a thin lens approximation for a 4× objective. This specific objective was used so that the simulated images would closely resemble the captured experimental images. For the simulation outlined in Figure 1a, the sources were distributed on a plane at the bottom of the scattering media. We refer to this arrangement of sources as “planar-bottom”.

This arrangement of sources (on a single plane) and tissue modeling with bulk optical parameters allows us to estimate a single source PSF for a given depth. We can then render a scene with multiple sources by convolving this estimated PSF with the source locations. After the scene was rendered, shot noise was added to account for the Poisson nature of light.

## 2 Computing the depth limit of bioluminescence

The depth limit in biological contexts is the point at which sources can no longer be resolved. This limit can be quantified in many ways. One common definition of this depth limit is the point at which two sources are no longer resolvable from one another (similar to the Rayleigh criterion) [22]. Alternatively, Pediredla *et al.* define the depth limit as the point at which the signal from the source equals the noise from the background [20]. Here, we take inspiration from Pediredla *et al.* and define depth limit as the farthest depth at which a point light source of a given brightness can be detected reliably (> 0.7 detection accuracy).

To quantify the reliability of detection, we use detection accuracy as a computational metric. Detection accuracy is defined as the 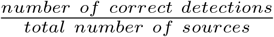 and is a quantifiable metric that describes how accurately we can detect bioluminescent sources within a scattering media. This can be thought of as the sensitivity of the system. To estimate the number of sources, we tested standard cell counting algorithms [9, 15]; however, these algorithms struggle in moderate scattering due to image blurring and signal attenuation. Compensating for signal attenuation by lowering the detection threshold increases the rate of false discoveries. For this reason, we developed an algorithm to detect sources in the presence of moderate and severe scattering.

Figure 2 illustrates the steps of the developed detection algorithm. Original images were rendered using the Monte Carlo framework described above. Our algorithm to detect sources consists of the following steps. Images were first blurred using a 2D Gaussian smoothing kernel with a standard deviation of 2 or 4 (chosen empirically) to reduce spurious detections resulting from shot noise. We then employed a local background subtraction step based on Bradley’s method [3] to overcome local fluctuations in the background due to the random distribution of sources. We performed this background subtraction on a per-pixel basis by subtracting from each pixel the average intensity of the neighboring pixels. This procedure generated the local background-subtracted image shown in the 3rd column of Figure 2. We then specified a sensitivity (threshold) to classify source and background pixels. Source pixels were then set to 1, and background pixels were set to 0 to create a binary image of source locations.

**Figure 2.**
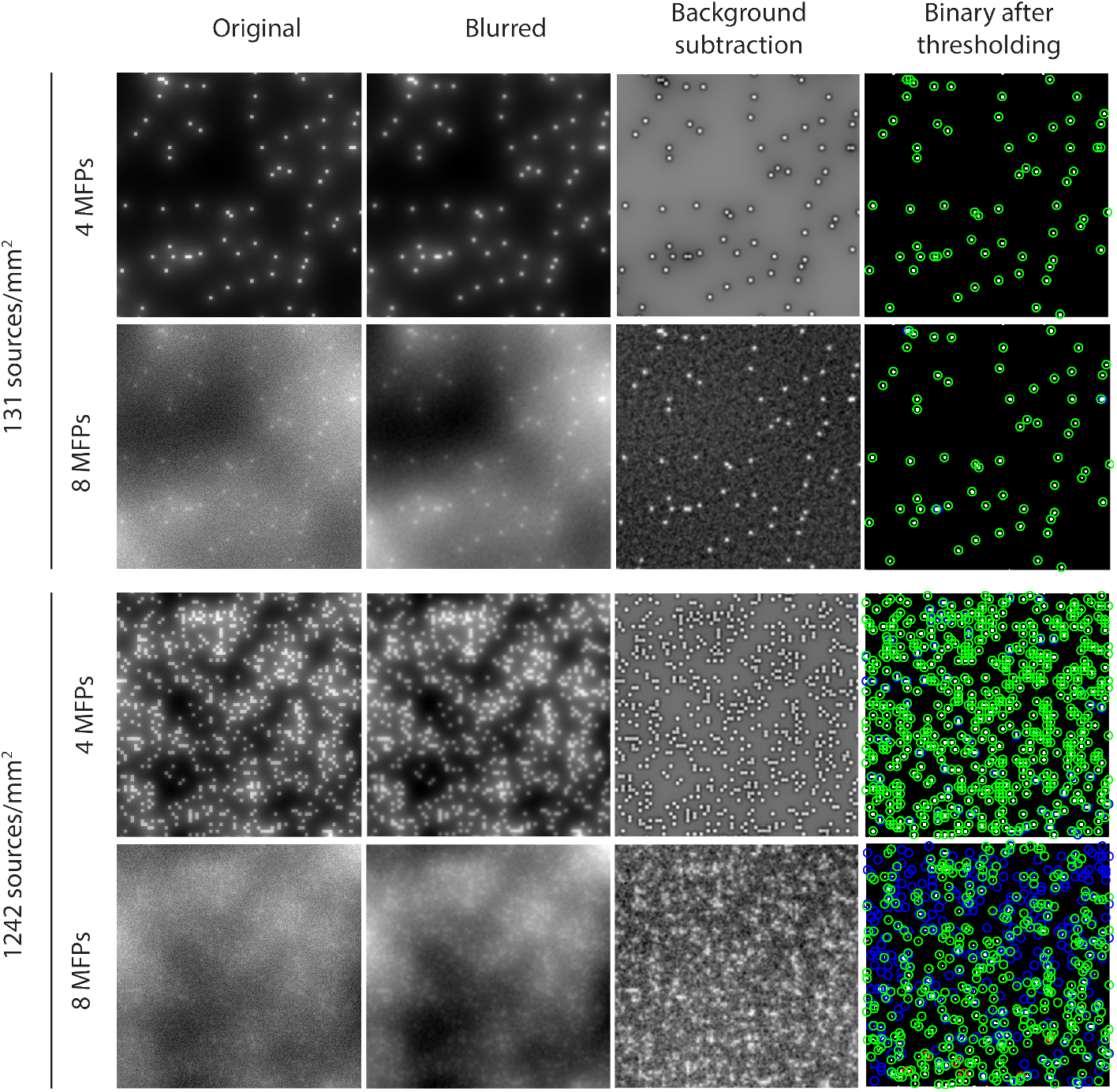
Detecting bioluminescent sources in captured images at different source densities and optical depths. Columns correspond to the steps taken by the detection algorithm going from the original image to the binary image after thresholding. The background subtraction step was based on Bradley’s method [3], which finds the background of the local region surrounding each pixel. Green, blue, and red circles represent correct, missed, and incorrect detections, respectively. Different source densities (131 and 1242 sources/mm^2^) and optical depths (4 and 8 MFPs) illustrate how the detection algorithm performs for different conditions. Detection accuracies from top to bottom are: 1.00, 0.97, 0.91, and 0.45. False discovery rates from top to bottom are: 0.00, 0.00, 0.00, and 0.05.

Detected source positions were derived from the centroids of binary source clusters. If a position was less than an empirically chosen distance (*20μm*) to any ground truth source location, the source was classified as a correct detection (green circles in *4^th^* column of Figure 2).

Detections outside this distance for all ground truth not already accounted for were registered as false discoveries (red circles in 4^*th*^ column of Figure 2). Ground truth locations not detected counted as missed detections (blue circles in 4^th^ column of Figure 2). Combinations of local neighborhood windows and sensitivities were tested for a given choice of system parameters (source density, emission rate) to realize the highest detection accuracy for a fixed false discovery rate. The window and sensitivity corresponding to the highest detection accuracy given a false discovery rate below the specified rate were used for the imaging parameters (optical depth, emission rate, and source density).

The rows of Figure 2 correspond to different source densities and optical depths. For all scenarios, the emission rate was 10^10^ photons/source. The detection accuracies corresponding to each row were 1.00, 0.97, 0.91, and 0.45 (read from top to bottom). The false discovery rates were all 0 except for the last row (1242 sources/mm^2^ and 8 MFPs), which was 0.05. Even in the presence of substantial scattering (third row of Figure 2), the detection algorithm performed well. However, at a higher source density (fourth row of Figure 2), the detection algorithm struggles to separate adjacent sources, resulting in lower detection accuracy.

Figure 1b demonstrates using detection accuracy to determine the depth limit. The bioluminescence model rendered images of sources located behind scattering media. These simulations used *l_s_* = 90*μm*, an MFP length matching the experimental MFP length for the brain phantoms generated for our experiments. Also, 90 *μm* is similar to the MFP length of green light in mouse brain tissue [20]. The simulated bioluminescent source density was 131 sources/mm^2^, and the emission rate was 10^10^ photons/source. The images shown in Figure 1b correspond to sources at a depth of 4, 8, and 10 MFPs. As the scattering depth increases, image quality degrades, and as a result, the detection accuracy starts to fall, as shown in the plot in Figure 1. After 8 MFPs, the detection accuracy falls considerably until no sources are distinguishable (10 MFPs). 8 MFPs is the depth limit of the system as it is the last thickness in which sources were detected reliably (detection accuracy of 0.7).

## 3 Characterizing bioluminescence imaging and experimental validation

Two primary factors affect imaging depth limit for a given MFP and imaging system: (a) brightness of the bioluminescent sources and (b) density of source distribution (which contributes to background signal level). Here, we investigate the role that these two factors play in the achievable depth limit using bioluminescence.

### 3.1 Emission rate and source density characterization

To investigate the impact of emission rate and source density on the depth limit of bioluminescence, we rendered images with varying photon rate per source and number of sources per simulation. For this parameter characterization, the same source arrangement was used as in Figure 1a (planar-bottom). Emission rates varied between 10^7^ and 10^11^ photons/source, and source densities varied between 10^1^ and 10^3^ sources/mm^2^. Figure 3a shows the results of this characterization. The colors in the heatmap correspond to the computed depth limit based on detection accuracies for each pair of emission rate and source density.

**Figure 3.**
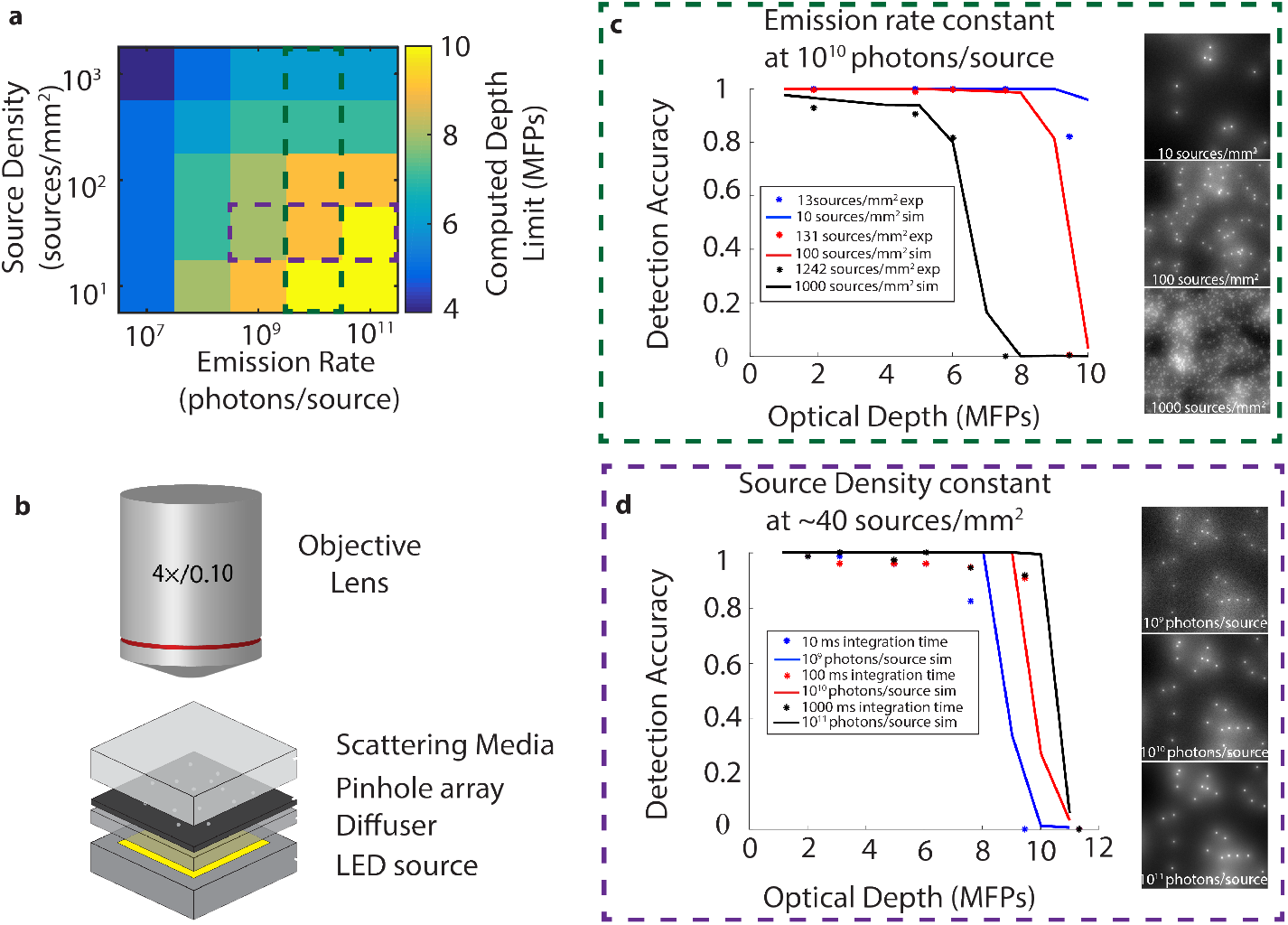
Depth limit of bioluminescence in scattering media as it changes with emission rate and source density, corroborated with experimental analog. (**a**) displays the computed depth limit as it varies with emission rate and source density. Computed depth limit determined to be the farthest depth that has a detection accuracy greater than 0.7. Green and purple boxed sections correspond to **c** and **d**, respectively. (**b**) shows the diagram of the experimental imaging setup. The LED source, diffuser, and pinhole array work together as the bioluminescent source analogy. (**c**) shows experimental results (asterisks) and simulated results (lines) for different source densities. Experimental and simulated results match fairly well for all three densities. The emission rate was kept constant and was 10^10^ photons/source for the simulation and 100 ms exposure time for experimental results. (**d**) shows experimental results (asterisks) and simulated results (lines) for different emission rates. Experimental and simulated results match fairly well for all three densities. Source density was kept constant 48 sources/mm^2^ for both simulation and experimental results.

As expected, a large emission rate and small source density yielded a higher computed depth limit. Conversely, a low emission rate and large source density yielded a smaller depth limit. Interestingly, many of the scenarios with 10^7^ photons/source had a small depth limit (5-6 MFPs). This is explained by the lack of signal arriving at the detector from these low emitting sources. For the rest of the pairs of parameters, there seemed to be a gradual increase in depth limit starting from low emission and large source density to large emission and low source density with a maximum computed depth limit of 10 MFPs.

### 3.2 Experimental bioluminescence analog for “planar-bottom” simulations

An experimental bioluminescence analog was designed in the planar-bottom source arrangement to corroborate findings from the simulations. A bioluminescent analog, rather than bioluminescent particles, was used to allow strong emitting sources and easy control of source density. The imaging setup is shown in Figure 3b. A diffuser was placed on an LED array to get a good angular distribution of light. A pinhole mask was placed on the LED array, restricting light to the 10 *μm* diameter pinholes (representing sources). PDMS tissue phantoms separated the pinhole sources from the imaging system to emulate imaging in tissue. Different integration times were used to change how many photons were imaged per source, and different densities of pinholes were used to achieve different densities of sources. A Blackfly Pointgrey camera was connected to an inverted microscope with a 4× objective to capture images.

The following are the steps carried out to create the PDMS phantoms. Polystyrene beads (0.6 mL) were added to isopropyl alcohol (0.6 mL). The solution of beads and

IPA were vortexed thoroughly for uniformity then added to 4 g of polydimethylsiloxane (PDMS). Again, the solution was mixed thoroughly for uniformity. Elastomer (0.4 grams) was added to the solution, and the solution was mixed thoroughly. The solution was left in the degassing chamber for 15 minutes or until all the bubbles evaporated from the solution. The solution was spun onto a glass slide to achieve the desired thickness (dependent on RPM). The wafer with the PDMS was baked in an oven at 37°C for 30 minutes. The tops of two phantoms were cleaned in the plasma chamber and then adhered to one another to make thicker phantoms. The stack of phantoms was put in the oven to bake for 30 minutes.

Figures 3c,d show experimental and simulated results plotted on the same axes. In Figure 3c, the simulation was kept at a constant emission rate of 10^10^ photons/source, and experiments were kept at a constant 100 ms exposure time. The three plots correspond to different source densities: 13 sources/mm^2^ (blue), 131 sources/mm^2^ (red), and 1242 sources/mm^2^ (black). The simulated and experimental detection accuracies follow similar trends and have a steep falloff around the same depth for the different source densities. In Figure 3d, source density was kept at a constant 48 sources/mm^2^ for both simulated and experimental results. While the direct relationship between simulated photons/source and experimental exposure time may be unclear, the results for 10 ms exposure time align well with the plot of 10^9^ photons/source (blue). Increasing the exposure time by 1 or 2 orders of magnitude should also increase the number of photons arriving at the sensor by the respective order of magnitude. Simulating this with 10^10^ (red) and 10^11^ (black) photons/source showed good agreement with 100 ms (red) and 1000 ms (black) integration times, respectively. The computed depth limits for Figure 3c,d are highlighted in Figure 3a with dashed boxes. Across all source densities and emission rates tested, experimental and simulated results agree, supporting the use of the simulation to compute the depth limit for bioluminescence imaging.

### 3.3 Feasibility of emission rates

Because the requirement for 10^7^ to 10^10^ photons per source is a potential limitation for bioluminescent imaging in scattering tissue, we investigated the feasibility of generating this signal magnitude in neurons. Currently, the brightest bioluminescent labels, such as the “eNanoLantern” series of FP-NanoLuc fusions [10,29], produce on the order of 10 photons/s per luciferase molecule. Thus, to achieve an emission rate of 10^7^ photons/s per neuron would require the soma to contain 10^6^ molecules of bioluminescent source, which corresponds to a cytosolic concentration of 2 *nM*. For comparison, fluorescent proteins expressed as unfused cytoplasmic markers typically generate protein concentrations in the micromolar range or higher [6, 21]. At similar expression levels of NanoLuc-based bioluminescent probe, a neuron could easily generate well over 10^10^ photons/s. We, therefore, conclude that the simulated and experimental imaging conditions shown here should already be possible with available bioluminescent probes.

### 3.4 3D imaging arrangement

All simulation and experimental results up to this point have had a planar-bottom source arrangement. It is of interest to consider the cases of sources fully embedded within the scattering media and sources not constrained to a single plane.

Figure 4 shows the impact of the planar-embedded and 3D-embedded source arrangements on detection accuracy. The left-most diagram of Figure 4 displays a 3D view and an X-Z projection of planar sources embedded within scattering media. The diagram to the right, 3D-embedded arrangement, illustrates sources with axial distribution across a depth of 300 *μm* (approximate size of V2&3 of the mouse brain [1,11]). In this comparison, the emission rate was fixed at 10^9^ photons/source. Planar arrangements have a source density of 13 sources/mm^2^, and the 3D arrangement has a source density of 100 sources/mm^3^. We chose a volumetric density of 100 sources/mm^3^ to closely match the amount of in-focus sources for a 4× objective as the planar source density. 3D simulations used a 75 *μm* MFP length (_l_s__) which directly reflects that of the mouse brain [20] (no experimental validation was performed).

**Figure 4.**
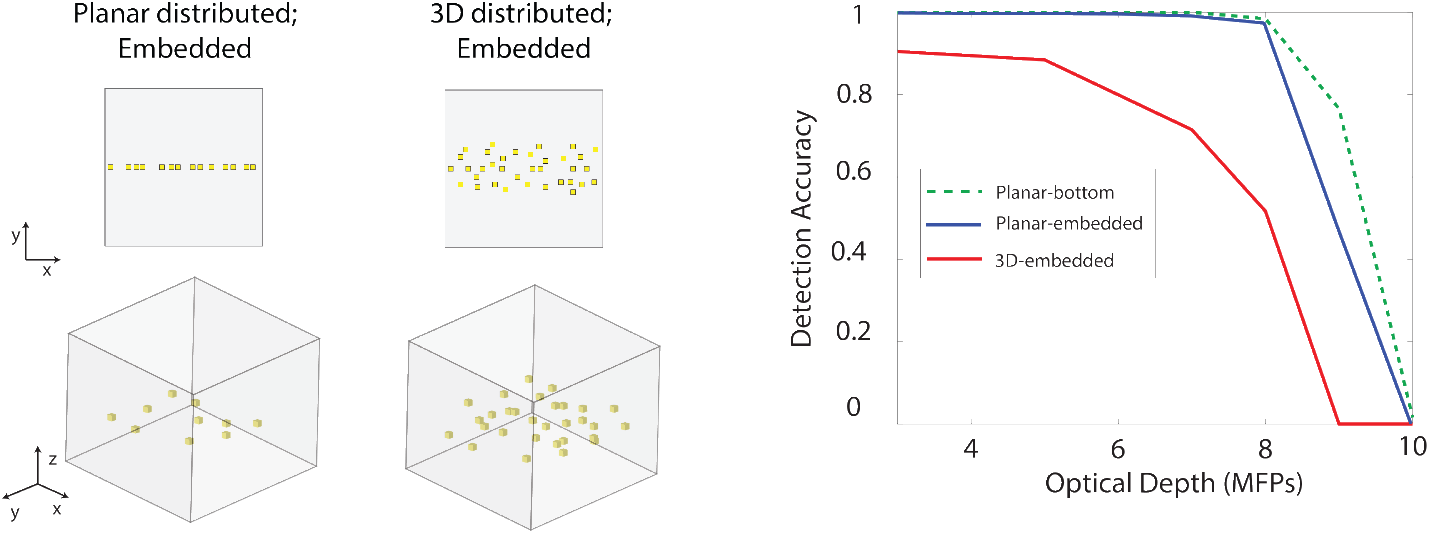
Depth limit of bioluminescence simulations with varying axial distributions. The two diagrams on the left show the planar-embedded and 3D-embedded source arrangements. The figure on the right shows how the detection accuracy changes with optical depth for the different source arrangements. The green dashed line, the blue line, and the red line represent the planar-bottom, planar-embedded, and 3D-embedded source geometries, respectively. Emission rate was kept constant at 10^9^ photons/source, and source density was 13 sources/mm^2^ for planar arrangements and 100 sources/mm^3^ for 3D arrangements.

In the case of a 3D-embedded source arrangement, the detection algorithm must be slightly modified to account for 3D positions. First, for a given source initialization, 7 different renderings, spaced 50 *μm* (depth of focus for 4× objective) apart, were performed. Planar detection of sources occurred the same as outlined above. The z-projection of the ground truth locations was used as the ground truth for each individual slice. Source locations that were detected in more than one axial slice were evaluated across slices to determine the axial position. The slice in which the full width at half maximum (FWHM) of the intensity profile was smallest was chosen. Acceptable axial distances from a detected source to a ground truth location were allowed to vary between 50*μm* (sources are shallow) to 100*μm* (sources are deep). This is in part due to the scattering blur outweighing the focusing blur deeper into the tissue. For axial detection, false discovery rates of 0.07 and below were tolerated.

The results of detecting planar-embedded and 3D-embedded source configurations are shown in the plot on the right of Figure 4. There is only a small difference in performance of the planar-embedded arrangement from previous planar-bottom simulations. The scattering medium is primarily forward scattering (*g* = 0.9), so most photons moving away from the detector will never end up reaching the detector. A depth limit of 8 MFPs is similarly achieved. On the other hand, having out-of-plane sources introduced blur, impeding the detection of sources at other planes (depth limit computed is 7 MFPs). The optical depths shown in the plot correspond to the axial center of the source distribution. The initial offset in the 3D-embedded plot is in part due to some sources being 1-2 MFPs deeper within the media.

## Conclusion

Through simulation and experimental verification, we have shown that bioluminescence microscopy is capable of imaging through 5-10 MFPs of scattering media. We have also demonstrated that the necessary signal intensity is currently possible with the brightest existing bioluminescent probes. This relatively deep imaging capability (compared to epifluorescence) combined with reduced phototoxicity, no photobleaching, and simple imaging hardware makes bioluminescence microscopy a valuable technology for *in vivo* imaging applications.

## Acknowledgments

This work was supported by a training fellowship from the Gulf Coast Consortia, on the IGERT: Neuroengineering from Cells to Systems, National Science Foundation (NSF) 1250104, grant R01GM121944, National Institutes of Health (NIH), and grant NeuroNex 1707352, NSF.

The authors would like to thank Adithya Pediredla for advise on the Monte Carlo simulation and potential avenues of improvement. The authors would also like to thank Vivek Boominathan for advice on improvements to the detection algorithm, as well as Christopher I. Moore and Caleb T. Kemere for useful discussions.

